# Presence of Ecuador in the Web of Science from open access in post-pandemic period 2019-2021: A multivariate analysis

**DOI:** 10.1101/2024.03.18.585617

**Authors:** Patricio Álvarez Muñoz, Fernando Erasmo Pacheco Olea, Dennis Alfredo Peralta Gamboa, Aviles Valenzuela Angelo Marcos

## Abstract

The study analyzed the presence of Ecuador in the post-pandemic Web of Science (WoS) from a multivariate dynamic, highlighting a significant transformation in scientific communication towards open access (OA). The methodology integrated multivariate statistical techniques, using R and specialized tools, to process 9,085 relevant articles. Between 2019 and 2021, 52% of Ecuadorian publications in WoS were open access. This movement reflects a commitment to quality research and accessibility, evidencing an increase in publications and citations under OA. The research highlights the importance of establishing robust national OA policies in Ecuador to promote scientific collaboration and an equitable distribution of knowledge. The dynamic biplot showed how universities in the post-pandemic period had stable and erratic behaviors towards open access. Of all the universities, the State University of Cuenca stands out in terms of quantity of production and migration to open access.

## Introduction

The COVID-19 pandemic has represented a turning point in contemporary history, profoundly impacting all sectors, including academia. In this context, scholarly communication has undergone a significant transformation, particularly in the field of open access (OA). This study focuses on Ecuador’s presence in the Web of Science (WoS) from open access in the post-pandemic period 2019-2021, exploring how the global crisis has accelerated the adoption of open access practices and influenced the country’s scientific production [1], [2, pp. 2006-2015].

With the choice of WoS as the main data source, given its recognized integrity and coverage in the global scientific literature [3], [4], the study embarked on a comprehensive analysis. Through a meticulous search strategy, 24,502 articles were initially identified, which were then reduced to 18,332 after a purification process, finally focusing on 9,085 publications from the post-pandemic period.

The R programming language was used, complemented by specialized tools for various functions, from data manipulation to advanced visualization [5]. This approach allowed a careful evaluation of the variables using the Bonferroni test [6], with a confidence level of 95%, in addition the use of the representation of the dataset using an HJ-Biplot allows a visualization and understanding of the migratory dynamics of the variables.

This change or migration towards more open models of higher quality research allows universities to have greater representation in the best databases in the world and in turn a higher level of citation and accessibility, despite the absence of specific regulations that promote open access in Ecuador. The study offers a clear vision of the evolution and impact of Ecuadorian scientific production in the global scenario, highlighting the influence of open access on scientific collaboration and the equitable distribution of knowledge [7], [8].

### State of the Art

The COVID-19 pandemic has catalyzed significant changes in scholarly communication, driving the open access (OA) movement. This change has been reflected in the growing adoption of OA policies by academic institutions and publishers worldwide, with a focus on the democratization of knowledge [9].

In Ecuador, the transition towards OA has been remarkable, especially in the context of the global health crisis. Universities and research centers have begun to adopt OA practices, resulting in an increase in publications on accessible platforms [10], [11, pp. 2010-2021].

The Web of Science, as the main repository of scientific literature, has registered an increase in contributions from Ecuadorian authors during and after the pandemic. This increase is indicative of the growing visibility and international impact of Ecuadorian research [12].

Despite progress, there are challenges to comprehensive OA adoption in Ecuador, including a lack of standardized policies and funding to cover article processing fees (APCs). However, these barriers also present opportunities for developing sustainable OA models that align with local needs and capabilities [13], [14]. The future of OA in Ecuador, especially in the post-COVID-19 landscape, looks promising. Current trends are expected to lead to increased international scientific collaboration and greater inclusion of Ecuadorian research in global discussions [15, pp. 2016-2020]. This study aims to analyze the presence of Ecuador in WoS from an open access perspective, using statistical tests, data visualization and a PCA Biplot representation to show university trends.

## Materials and methods

We collected data from the source: Web Of Science with all its collections such as: SCIE, SSCI, AHCI, ESCI, CPCI, BKCI. The data were retrieved until August 17, 2023, the search was performed through the following equation: ‘CU=(ECUADOR) AND PY=(2011-2021) AND DT=(Article)’.

The analysis was performed from 2019-2021; multivariate statistical, bibliometric and data visualization techniques were used with the objective of analyzing the presence of Ecuador in the Web of Science (WoS) database. With an exclusive focus on the context of open access.

First, the scientific information database Web of Science (WoS) was chosen due to its recognized integrity and coverage of the global scientific literature and rigorous indexing criteria in the academic and research sphere [16].

Keywords and descriptors related to Ecuador and open access were used, resulting in 24,502 articles, to consolidate in a single data the articles, 25 batches of 1000 articles were downloaded, then using a Linux virtual machine (Kali Linux), the batches were combined using the cat command.

Using data wrangling techniques, the data was read and debugged. Once consolidated, any variables with repeated identifiers were removed, ensuring the accuracy and integrity of the data set. As a result of this process, the total number of articles was reduced from 24502 to 18332 (we will use the term observations in order to apply statistical analysis techniques to our data). For the study, we statified the data by taking from the year 2019 to the year 2021. This leaves a total of 9085 data to be analyzed.

### Analysis

For data analysis, the R programming language was used, complemented with various specialized tools:

dplyr: Facilitates data manipulation and transformation.
stringr: Helps in the management and manipulation of text strings.
openxlsx: Allows reading and writing Excel files.
ggplot2: Visualization tool for creating complex graphs.
reshape2: Used to restructure and aggregate data.
corrplot: Generates correlation graphs.
factoextra: Assists in extracting and visualizing results from various data analyses.
gridExtra: Provides functions to combine multiple ‘ggplot’ plots. nbclust, cluster: Tools to determine the optimal number of clusters and perform cluster analysis.
cowplot: Enhance and customize plots.
pandas (Python): Data analysis library that provides flexible data structures.

These tools were used to segment and categorize Ecuador’s universities according to their nature (public and private) and the type of access to their publications (open or subscription). Through this approach, it was possible to evaluate the distribution of citations and their annual average, and to highlight publications under open access models [17].

For an effective graphical representation of the results, advanced data visualization techniques were used. On the statistical side, a detailed analysis of qualitative and quantitative variables divided into factors or groups was carried out, implementing a Bonferroni test to identify significant differences in the number of citations between groups [18].

To obtain a better representation of Universities and publication sources (Open Access and Subscription) this dataset was represented in a dynamic HJ-Biplot in the period 2019-2021 (Yamashita & Mayekawa, 2015), the aim is to obtain the quality of the data representations with the following information:

1. Acute angles represent strong direct correlation.
2. Obtuse angles represent an inverse correlation.
3. Right angles represent independence.
4. Distance between observations (universities) indicates similarity.
5. The length of the vectors approximate the standard deviation of the vectors.

## Results and discussion

The scientific production and the number of citations of the publications of Ecuadorian universities in WoS has increased steadily in the post-pandemic period, as shown in Figure 1.

**Fig 1.**
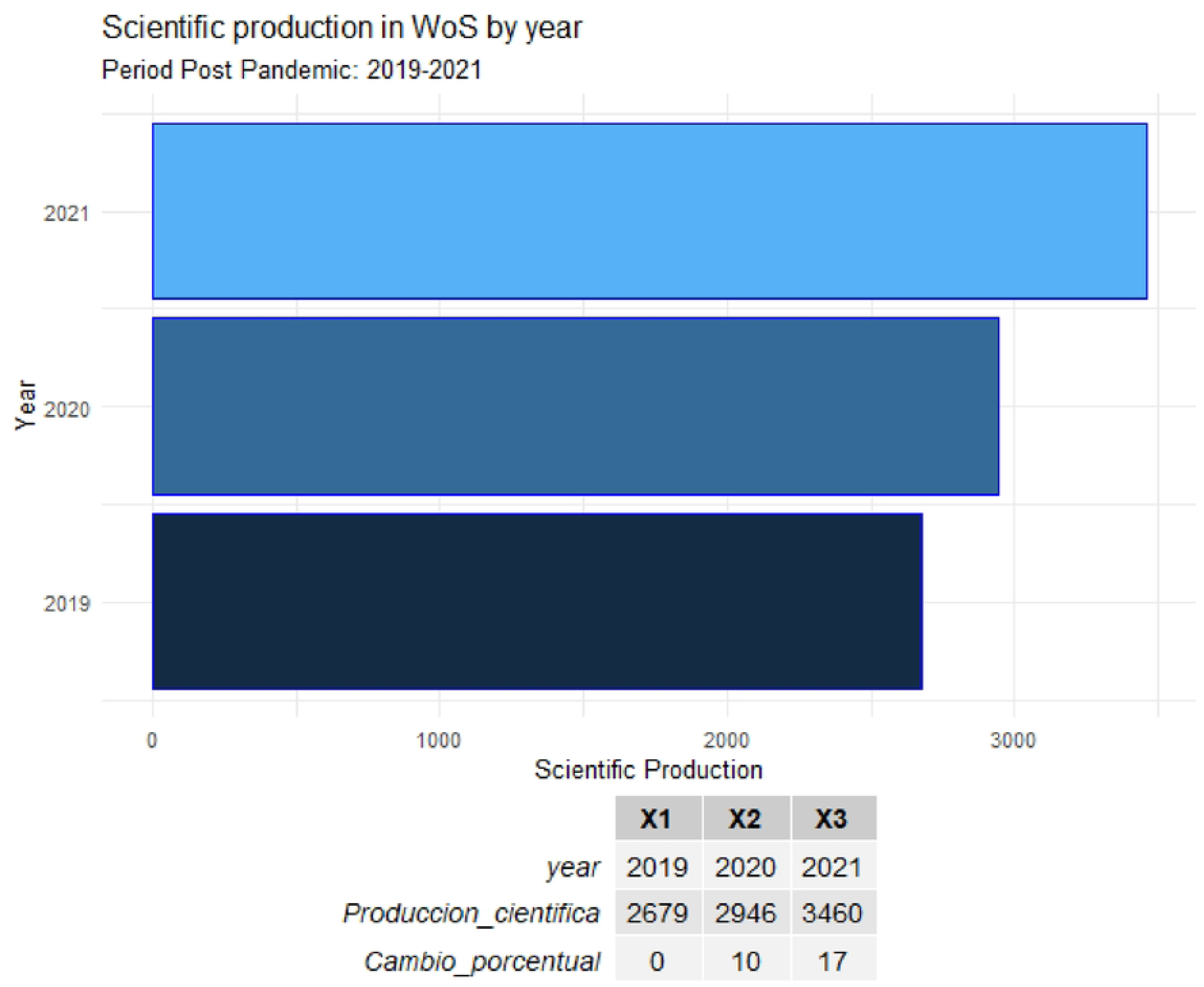
Scientific production and Number of citations of production in WoS by year. *Note*. Source: Web of science consulted as of August 17, 2023. Elaboration: Authors

Regarding Ecuador’s trend in open access, 4759 publications belong to Open Access journals out of a total of 9085 scientific articles. This implies that 52% of Ecuadorian publications in WoS in the period studied are under this modality, as shown in Figure 2.

**Fig 2.**
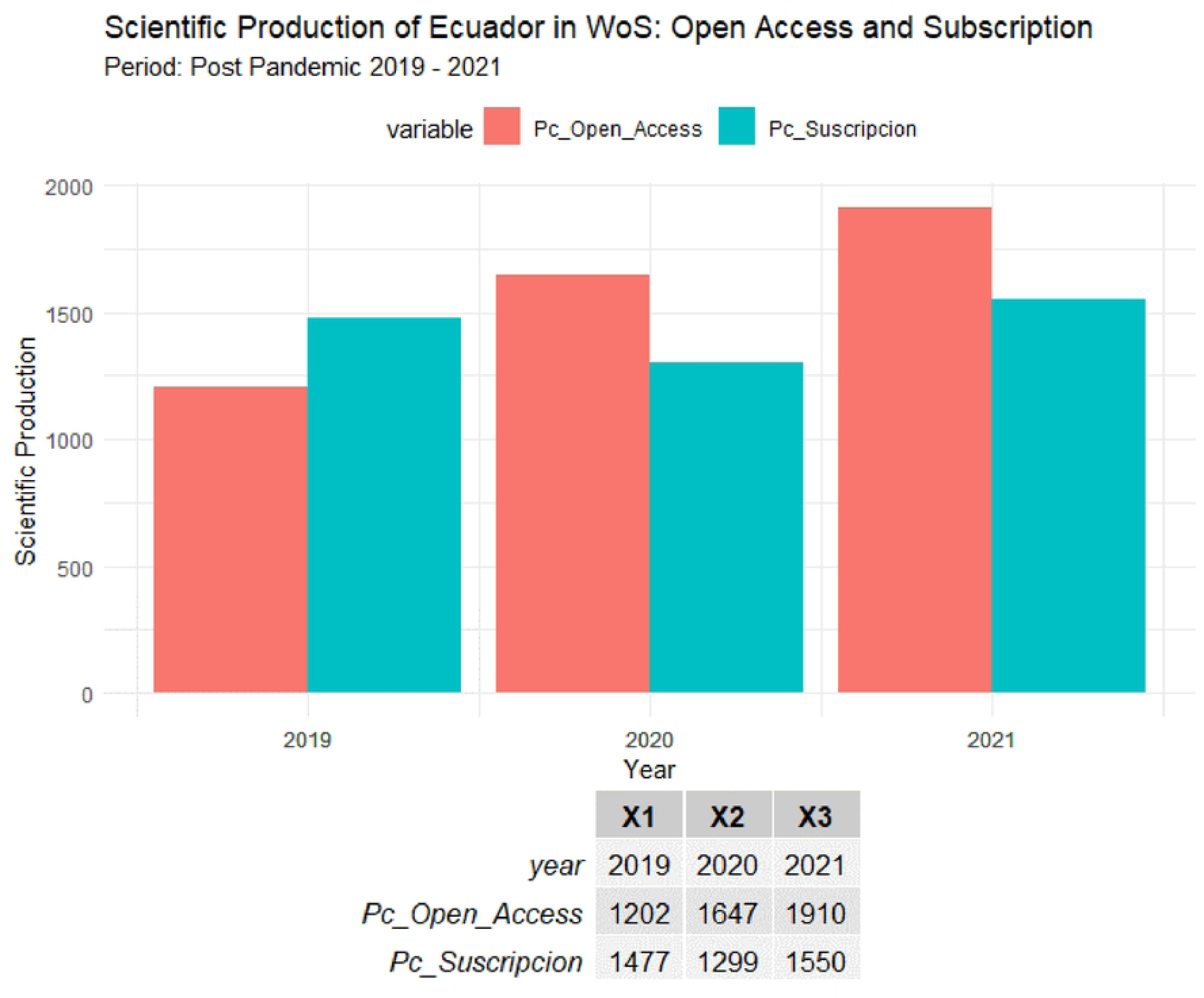
Scientific production of Ecuador in WoS: Open Access and Subscription. *Note*. Source: Web of science consulted as of August 17, 2023. Elaboration: Authors

Until 2019 publications in subscription journals outnumbered those in open access journals, this has changed as of 2020. Regarding the number of citations of scientific publications, Figure 3 shows that publications in open access journals receive more citations than those published in subscription journals.

**Fig 3.**
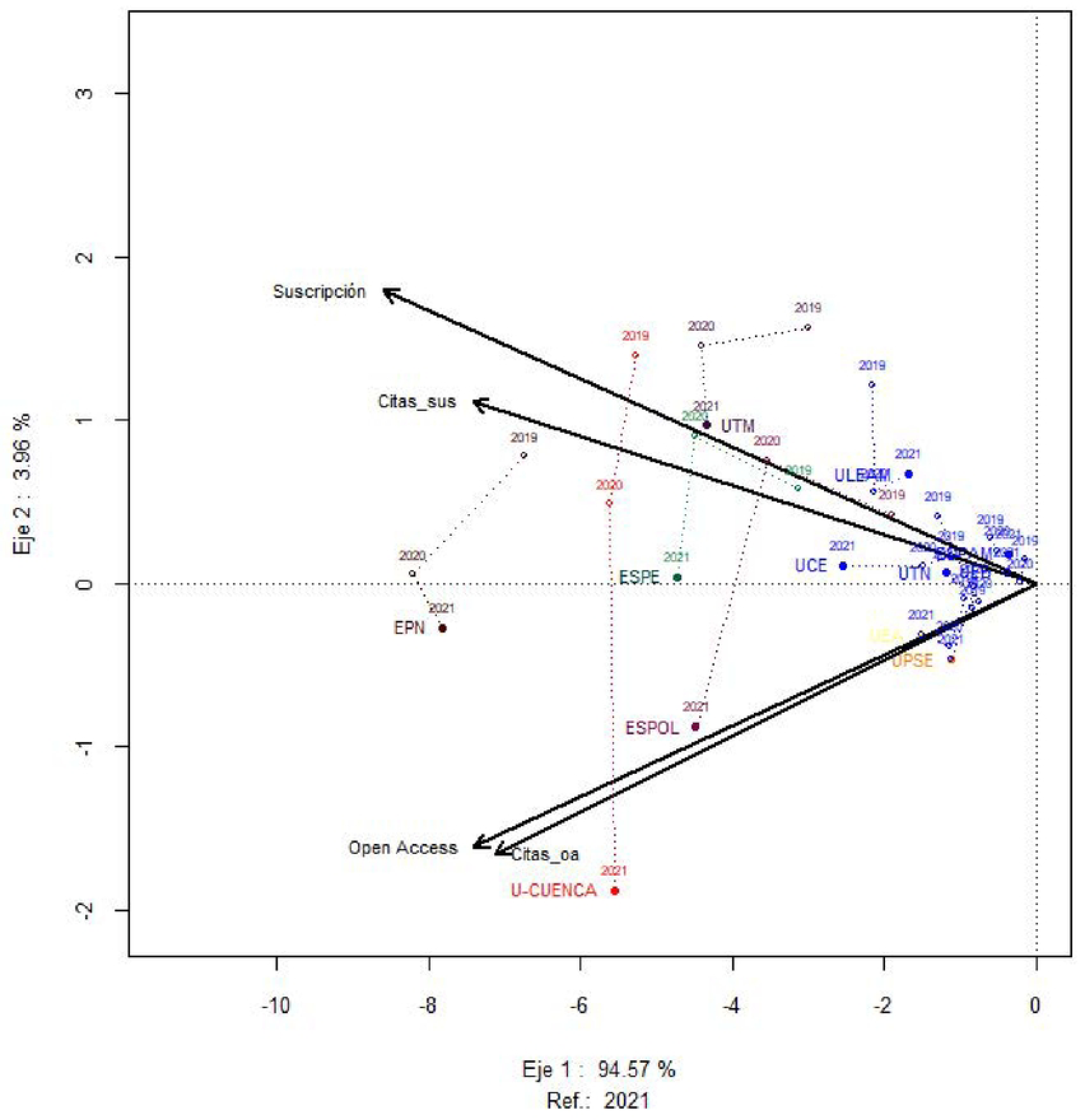
HJ-Biplot Multivariate dynamics of public universities. *Note*. Source: Web of science consulted as of August 17, 2023. Elaboration: Authors

These differences in citations were tested using a Bonferroni test at 95% confidence. The results showed to be significant (p=0), thus confirming that an article published in open access journals receives on average more citations than articles published in subscription journals.

Private institutions register 3905 scientific publications in WoS, while public universities register 3846 publications in the same database. 1,334 scientific publications are the product of collaboration between public and private universities. As shown in Table 1, these figures represent 43%, 42% and 15% of the total, respectively.

**Table 1.** Scientific publications of Ecuador in WoS period 2019-2021.

| Year | Pc_public | Pc_private | Pc_colaboration |
| --- | --- | --- | --- |
| 2019 | 1259 | 1045 | 375 |
| 2020 | 1351 | 1178 | 417 |
| 2021 | 1236 | 1682 | 542 |
| **Total** | 3846 (42%) | 3905 (43%) | 1334 (15%) |

Web of science accessed as of August 17, 2023.
<sup>a</sup>Elaboration: Authors using R .

Analyzing the trend of publications in open access, Table 2 shows that, of the 3846 articles from Ecuadorian public universities, 2018 were published in open access journals and 1828 in subscription journals. Of the 3905 research papers from private universities, 1943 appeared in open access journals and 1962 in subscription journals. Finally, of the 1334 collaborative research studies, 798 were published in open access journals and 536 in subscription journals.

**Table 2.** Open access scientific production private university vs public university.

| **Year** | **Public production of access open access** | **Open access private production** | **Open access collaborative production** | **Public subscription production** | **Private subscription production** | **Production private subscription partnership** |
| --- | --- | --- | --- | --- | --- | --- |
| **2019** | 568 | 493 | 141 | 691 | 552 | 234 |
| **2020** | 742 | 637 | 268 | 609 | 541 | 149 |
| **2021** | 708 | 813 | 389 | 528 | 869 | 153 |
| **Total** | 2018 | 1943 | 798 | 1828 | 1962 | 536 |

Web of science accessed as of August 17, 2023.
<sup>a</sup>Elaboration: Authors using R .

Research published in open access journals constitutes 52.4%, 49.7% and 59.8% of the total publications of public, private and collaborative universities, respectively. This behavior demonstrates that open access is increasingly present in Ecuador’s collaborative research ecosystems.

Figure 3 shows an HJ-biplot generated from a multivariate statistical analysis with a representation quality of 950 for the observations (Universities). It visualizes the relationship between different universities and the quantity of their publications in two categories, Open Access and Subscription, over several years (2019, 2020 and 2021). The orientation and length of the vectors indicate how each variable contributes to the variation in the data and the relationship between them. The proximity of the points to each vector suggests the magnitude of each variable for that observation

The first dimension, Axis 1, explains 94.57% of the variance, indicating that most of the information on the differences between universities is along this axis, so the main variability is in the number of publications and citations, whether Open Access or Subscription.

As expected there is a high direct correlation between the number of citations and articles in Open Access and subscription journals. Universities that are located at the top of Axis 1 have a better representation for concentrating their scientific publications in Open Access journals and those that are drawn at the bottom of Axis 1 have a better representation for publishing their research in subscription journals.

Publications in subscription journals have a positive correlation with most universities over the years, while publications in open access journals have a less significant relationship with public universities. The oldest universities in Ecuador such as the Escuela Superior Politécnica del Litoral (ESPOL), the Escuela Politécnica Nacional (EPN) and the Universidad Estatal de Cuenca (U-CUENCA) show a remarkable change in publication sources in the post pandemic period, initially in 2019 these universities were best represented by concentrating their publications in subscription journals and by the end of 2021 a latent migratory trend to open access is observed.

Universities located near the origin of coordinates such as Universidad Técnica del Norte (UTN) and Universidad Central del Ecuador (UCE) suggest a lower number of publications and citations in both categories compared to other public universities in Ecuador. The Polytechnic University of Santa Elena (UPSE) and the Agrarian University of Ecuador are located in an atypical case, since they are the ones that have all their publications tending to open Access.

The study highlights the preferences of Ecuadorian researchers when publishing their articles under the open Access model. Figure 4 illustrates a clear tendency among academics from Ecuadorian universities, both from the public and private sectors, to choose which open access routes they choose for the publication of their research in WoS. The evidence according to the data reflects that the Green option (which allows authors to archive versions of their work in specific repositories), leads with 2425 publications. This figure exceeds the 1955 publications that chose the Gold route (which involves direct publication in open access journals), the 200 that chose the Bronze route, and the 179 that chose the Hybrid route (where certain articles in traditional journals are made accessible to the public), demonstrating that these routes are not desired by Ecuadorian researchers to disseminate their findings.

**Fig 4.**
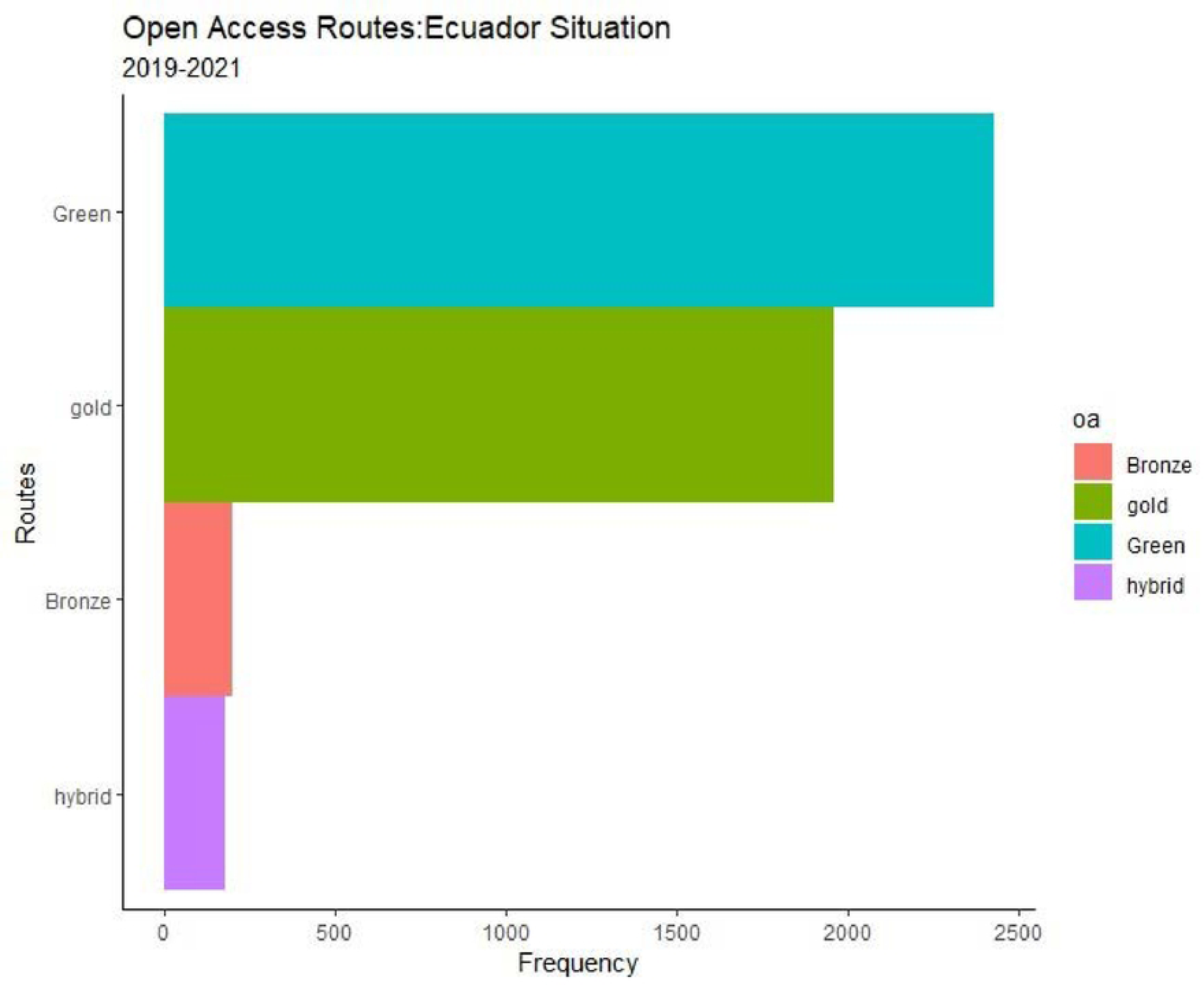
Open Access scientific production of Ecuador in the Web of Science. *Note*. Source: Web of science consulted as of August 17, 2023. Elaboration: Authors

## Discussion

This study reveals a significant change in Ecuador’s scientific publication strategy in the context of open access, especially during and after the COVID-19 pandemic. The adoption of open access in Ecuador, with 52% of publications in this format during the period studied, reflects a global trend towards greater transparency and accessibility in scientific communication [19]. However, unlike countries with more consolidated OA policies, as observed in the study by [20], Ecuador shows an emerging and developing approach, which underscores the importance of establishing more robust national OA policies.

When comparing the scientific production and citations of Ecuadorian universities in WoS in the period studied 2019-2021, it is observed that open access publications receive on average more citations than subscription publications, which is consistent with the findings of [21]. This pattern suggests that OA not only improves the visibility of research but also its impact, an aspect highlighted in the literature on science communication [22].

Collaboration between public and private universities in Ecuador, representing 15% of total publications, is remarkable but still lower than that observed in more collaborative contexts such as Europe, as reported by [23]. This indicates a potential for growth in inter-institutional collaboration in the country, with much more strength in the post-covid period.

In terms of the distribution of publications by type of access, there is evidence of a growing preference for open access journals in Ecuador, which is in line with the post-pandemic global trends observed by [24]. However, Ecuador still faces challenges in terms of comprehensive adoption of these practices, compared to more advanced countries in this aspect, as described in the research by [25].

Finally, while Ecuador shows progress in the adoption of open access and increasing presence in WoS, there is still room for improvement, especially in terms of national policies and inter-institutional collaboration, as institutions lack clear regulations that stimulate open access production.

The study is consistent with the current literature and highlights the need for a more proactive and systematic approach to OA in the field of scientific research in Ecuador.

## Conclusions

The analysis carried out has shown a significant transformation in scientific communication in Ecuador, especially marked by the adoption of open access (OA). This trend, which advocates the free availability of scientific literature, has profoundly altered the paradigm of publication and distribution of knowledge.

These data reflect a strategic adaptation to the post-pandemic global environment and a response to the growing demand for accessible and transparent scientific information. Despite financial challenges, reduced government spending on higher education and changing perceptions about the quality of publications, educational institutions in Ecuador have demonstrated an ability to maintain a robust and consistent scientific output, adapting to changes in the scientific publishing landscape.

Ecuador’s incorporation into the global OA landscape is a significant step towards the democratization of knowledge. However, to take full advantage of the benefits of OA, a more integrated strategy and sustained legal support at the national level is required. This implies not only encouraging OA publishing, but also adopting research and education policies that support the creation and dissemination of knowledge in an open and accessible manner, which is currently very incipient.

In conclusion, the study underscores the need for a holistic vision and approach to OA in Ecuador. The researchers have done their part, despite adverse circumstances.

## References

[1] S. Minniti, V. Santoro, y S. Belli, «Mapping the development of Open Access in Latin America and Caribbean countries. An analysis of Web of Science Core Collection and SciELO Citation Index (2005–2017)», Scientometrics, vol. 117, n.o 3, pp. 1905–1930, 2018, doi: 10.1007/s11192-018-2950-0.

[2] J. A. Castillo y M. A. Powell, «Analysis scientific production from Ecuador and the impacto of international collaboration in the period 2006-2015», Revista Espanola de Documentacion Cientifica, vol. 42, n.o 1, 2019, doi: 10.3989/redc.2019.1.1567.

[3] V. V. Macchi Silva, J. L. Duarte Ribeiro, G. R. Alvarez, y S. E. Caregnato, «Competence-Based Management Research in the Web of Science and Scopus Databases: Scientific Production, Collaboration, and Impact», Publications, vol. 7, n.o 4, p. 60, dic. 2019, doi: 10.3390/publications7040060.

[4] A. H. Sawalha, D. H. Solomon, K. D. Allen, P. Katz, y E. Yelin, «Immediate Open Access: The Good, the Bad, and the Impact on Academic Society Publishing», ACR Open Rheumatology, vol. 5, n.o 6, pp. 308–309, 2023, doi: 10.1002/acr2.11547.

[5] M. M. Julkowska et al., «MV app-multivariate analysis application for streamlined data analysis and curation», Plant Physiology, vol. 180, n.o 3, pp. 1261–1276, 2019, doi: 10.1104/pp.19.00235.

[6] T. D’Isanto, G. Altavilla, G. Esposito, G. Raiola, y F. D’Elia, «Physical activity and sports sciences field in Italian scientific research products and its distinct composition in biomedicine, exercise and sports sciences and pedagogy domains», Sport Sciences for Health, vol. 19, n.o 3, pp. 987-991, 2023, doi: 10.1007/s11332-023-01045-z.

[7] M. G. Claudio-González y A. Villarroya, «Challenges of publishing open access journals», Profesional de la Informacion, vol. 24, n.o 5, pp. 517–525, 2015, doi: 10.3145/epi.2015.sep.02.

[8] P. Yuanyuan, H. Jinxia, C. Xuefei, y Z. Zhanyi, «Open Access Journals’ Development in the Open Science Process(2017-2020)», Journal of Library and Information Science in Agriculture, vol. 32, n.o 12, pp. 29–40, 2020, doi: 10.13998/j.cnki.issn1002-1248.2020.12.20-0949.

[9] D. Torres-Salinas, «Daily growth rate of scientific production on covid-19. Analysis in databases and open access repositories», Profesional de la Informacion, vol. 29, n.o 2, 2020, doi: 10.3145/epi.2020.mar.15.

[10] C. M. Beitl, «Navigating Over Space and Time: Fishing Effort Allocation and the Development of Customary Norms in an Open-Access Mangrove Estuary in Ecuador», Human Ecology, vol. 42, n.o 3, pp. 395-411, 2014, doi: 10.1007/s10745-014-9655-7.

[11] C. H. González-Parias, J. A. Londoño-Arias, y W. A. Giraldo-Mejía, «EVOLUTION OF SCIENTIFIC PRODUCTION IN LATIN AMERICA INDEXED IN SCOPUS 2010-2021», Bibliotecas, Anales de Investigacion, vol. 18, n.o 3, pp. 1–14, 2022.

[12] B. Probst, P. M. Lohmann, A. Kontoleon, y L. D. Anadón, «The impact of open access mandates on scientific research and technological development in the U.S.», iScience, vol. 26, n.o 10, 2023, doi: 10.1016/j.isci.2023.107740.

[13] S. Belli, R. Cardenas, M. Velez, A. Rivera, y V. Santoro, «Open science and open access, a scientific practice for sharing knowledge», presentado en CEUR Workshop Proceedings, 2019, pp. 156–167.

[14] N. A. Mazov y V. N. Gureyev, «Open Access Bibliographic Resources for Maintaining a Bibliographic Database of Research Organization», Scientific and Technical Information Processing, vol. 50, n.o 3, pp. 211-223, 2023, doi: 10.3103/S0147688223030115.

[15] Q. T. Eneida María, R. L. Felipe, C. M. Exio Isaac, y M. I. Juan Carlos, «Scientific production on social responsibility in the social economy according to Scopus, period 2016-2020», Revista de Ciencias Sociales, vol. 28, n.o 2, pp. 258–275, 2022, doi: 10.31876/rcs.v28i2.37937.

[16] C. Birkle, D. A. Pendlebury, J. Schnell, y J. Adams, «Web of science as a data source for research on scientific and scholarly activity», Quantitative Science Studies, vol. 1, n.o 1, pp. 363–376, 2020, doi: 10.1162/qss_a_00018.

[17] X. Bosch, «A reflection on open-access, citation counts, and the future of scientific publishing», Archivum Immunologiae et Therapiae Experimentalis, vol. 57, n.o 2, pp. 91–93, 2009, doi: 10.1007/s00005-009-0016-y.

[18] B. Ljesević, Z. Martinović, M. Popović, y S. Jović, «[Visual vs. quantitative electroencephalographic analysis in patients with and without posttraumatic epilepsy].», Medicinski pregled, vol. 63, n.o 1-2, pp. 40–46, 2010, doi: 10.2298/MPNS1002040L.

[19] A. O. Rivero-Guerra, «The scientific production of nature-based tourism: bibliometric analysis of the Clarivate Analytics databases», Rev. Gen. Inf. Doc., vol. 31, n.o 1, pp. 461–493, 2021, doi: 10.5209/rgid.76973.

[20] S. Moradi y S. Abdi, «Open science-related policies in Europe», Science and Public Policy, vol. 50, n.o 3, pp. 521–530, 2023, doi: 10.1093/scipol/scac082.

[21] S. G. Akterian, «Towards open access scientific publishing», Biomedical Reviews, vol. 28, pp. 125–133, 2017, doi: 10.14748/bmr.v28.4459.

[22] D. de Filippo y J. Mañana-Rodriguez, «Open access policies and mandates and their practical implementation in Spanish public universities», presentado en 18th International Conference on Scientometrics and Informetrics, ISSI 2021, 2021, pp. 317–328.

[23] T. Piazzini, «Open access as a new paradigm. An inevitable evolution?», JLIS.it, vol. 11, n.o 3, pp. 99–109, 2020, doi: 10.4403/jlis.it-12631.

[24] G. F. Nane, N. Robinson-Garcia, F. van Schalkwyk, y D. Torres-Salinas, «COVID-19 and the scientific publishing system: growth, open access and scientific fields», Scientometrics, vol. 128, n.o 1, pp. 345-362, 2023, doi: 10.1007/s11192-022-04536-x.

[25] N. Chakravorty, C. S. Sharma, K. A. Molla, y J. K. Pattanaik, «Open Science: Challenges, Possible Solutions and the Way Forward», Proceedings of the Indian National Science Academy, vol. 88, n.o 3, pp. 456–471, 2022, doi: 10.1007/s43538-022-00104-2.

